# Differences in intake of high-fat high-sugar diet are related to variations in central dopamine in humans

**DOI:** 10.1101/839647

**Authors:** Hendrik Hartmann, Larissa K. Pauli, Lieneke K. Janssen, Sebastian Huhn, Uta Ceglarek, Annette Horstmann

## Abstract

Obesity is associated with alterations in dopaminergic transmission and cognitive function. Recent findings from rodent studies suggest that diets rich in saturated fat and refined sugars (HFS) induce changes in the dopamine system independent of excessive body weight. However, so far the impact of HFS on the human brain has not been investigated. Here, we compared the effect of dietary dopamine depletion on dopamine dependent cognitive tasks between two groups that differ in habitual intake of dietary fat and sugar. Specifically, we used a double-blind within-subject crossover design to compare the effect of acute phenylalanine/tyrosine depletion (APTD) on a reinforcement learning and a working memory task, in two groups that are on opposite ends of the spectrum of self-reported HFS intake (low vs. high intake: LFS vs. HFS group). We tested 31 healthy young women, who were matched for BMI (mostly normal weight to overweight) and IQ. Depletion of central dopamine reduced the working memory specific performance on the operation span task (OSPAN) in the LFS, but not in the HFS group (p = 0.023, r = 0.210). Learning from positive and negative reinforcement (probabilistic selection task: PST) was increased in both diet groups after dopamine depletion (p = 0.048, r = 0.144). As secondary exploratory research question we measured peripheral dopamine precursor availability (pDAP) at baseline as an estimate for central dopamine levels. The HFS group had a significantly higher pDAP at baseline compared to the LFS group (p = .048, r = −0.355). Our data provides first evidence that the intake of HFS is associated with changes in indirect measures of central dopamine levels in humans. The observed associations are independent of body weight status, suggesting that consumption of HFS might be associated with maladaptive behaviors contributing to the development of obesity.

## 1. Introduction

Over the last decades, obesity has become a global health burden, making research on the development and maintenance of obesity more relevant than ever. One of the main drivers of the rapid rise in obesity rates is the increased intake of food products containing high amounts of saturated fat and refined sugars^1^. The question is, what makes people eat beyond their caloric needs, despite negative consequences, such as getting uncomfortably full or the health risks associated with obesity?

Throughout their daily life people are constantly exposed to food advertisements and easily available food products. Such external food cues have the potential to enhance the motivation to obtain and consume food, even in a satiated state^2^. Recently, it has been shown that people with obesity outperform people with normal weight when learning and tracking the reward predicting value of cues associated with a food reward^3^. In addition, individuals with higher BMI compared to lower BMI (normal weight to obese) continue to respond to such food reward cues with the same intensity, despite their decreased motivation to consume the food rewards after devaluation^4^. In a meta-analysis, Garcia-Garcia and colleagues^5^ showed that people with obesity exhibit hyperactivation in reward-related brain areas and proposed that this enhanced focus on rewards may lead to compulsive-like behaviors. In addition to motivational aspects and behavioral control, obesity is associated with altered decision making and executive functions^6,7^. Adverse decision making might be explained by the inability to integrate negative feedback as shown by impaired reinforcement learning associated with obesity^8^. In a probabilistic reinforcement learning paradigm with monetary rewards people with obesity chose the correct option less frequently and gained lower overall payout than lean participants^9^. Coppin and colleagues^10^ report similar findings of impaired reinforcement learning in obesity, but also impairments of working memory in line with previous findings^11,12^. The observed alterations of cognitive processes linked to motivation and behavioral control may contribute to the maintenance of obesity and are thought to be due to alterations in central dopamine pathways (reviewed by^6^). Reinforcement learning and working memory both depend on action of dopamine in the striatum and prefrontal cortex (PFC), and optimal levels are crucial for proper functioning^13–15^.

Although alterations in the dopaminergic system have mainly been associated with body weight in humans^16–21^, studies in rodents suggest that it is, in fact, a diet high in saturated fat and refined sugar (HFS) that leads to the observed changes, independent of excessive weight: exposure to high-fat diets reduced dopamine receptor D2 protein expression levels^22^, affected dopamine synthesis^23,24^, and uptake of striatal dopamine in rodents^25,26^. Furthermore, overconsumption of specifically saturated dietary lipids, predominating in a typical western style diet, reduced dopamine receptor D1 signaling in rats independent of weight gain^27^. Mimicking the effects of hidden sugars in commercial foods and beverages, low-concentration sucrose solutions changed dopamine receptor D1 and D2 mRNA and protein expression in the striatum^28^. Further, a high-fat diet down-regulated the expression of striatal dopamine receptor D1 and D2 mRNA^29^. However, it is not clear whether the observed alterations of the dopaminergic system are directly caused by HFS or are compensatory adaptations in response to altered dopamine levels.

Taken together, HFS may thus be responsible for the observed differences in adaptive behavior that crucially rely on the neurotransmitter dopamine and that promote the overconsumption of such food products and obesity. However, translating findings from animal to human studies has to be done with great care due to the large knowledge gap between the fields^30^. Up to date a possible relationship between HFS and the dopamine system has not been investigated in humans. Here, we aim to find evidence that high (relative to low) dietary intake of saturated fat and free sugars is associated with alterations of central dopamine and dopamine-dependent cognition, particularly, reinforcement learning and working memory.

Because the synthesis of monoamine neurotransmitters in the brain depends on the availability of their amino acid precursors circulating in the blood (peripheral dopamine precursor availability: pDAP), central dopamine levels can be decreased by depleting its precursors tyrosine and phenylalanine relative to the other large neutral amino acids, which competitively share a carrier at the blood brain barrier^31^. To what extent exactly central dopamine levels are decreased by precursor depletion and whether this decrease is of similar magnitude between individuals is not known; higher concentrations of striatal dopamine or presynaptic dopamine synthesis capacity, as shown for women compared to men^32,33^, could serve as buffer for the effects of peripheral depletion. To uncover potential diet-related differences in central dopamine and dopamine-mediated cognition, we made use of an acute phenylalanine/tyrosine depletion (APTD) method, which attenuates dopamine synthesis and release in the striatum^34–36^ and impairs frontostriatal functional connectivity^37^. APTD has been shown to modulate reinforcement learning^38,39^ and executive functions like set-shifting and spatial working memory^37,40,41^. These cognitive processes require a certain level of dopamine for optimal performance. Either a decrease or increase in this level will lead to suboptimal performance, i.e. dopamine levels relate to cognitive performance in an inverted-u shaped manner^15^. As such, assessing reinforcement learning and working memory performance after reduction of central dopamine availability in two groups that differ markedly in their dietary intake of saturated fat and free sugars could reveal potential diet-related differences in the dopamine system. We assessed reinforcement learning using a probabilistic selection task^14^ and working memory using the operation span task^42^ after APTD and after a balanced amino acid condition in a within-subjects design. We hypothesized that (1) high relative to low dietary intake of saturated fat and free sugars – as assessed using the Dietary Fat and free Sugar Questionnaire (DFS)^43,44^ – is associated with altered performance on both tasks, and (2) that APTD may differentially affect performance of the two diet groups depending on diet associated differences in their dopaminergic system.

## 2. Methods

### 2.1. Participants

90 healthy female participants (age, 25.03 ± 3.61 years; BMI, 24.16 ± 5.72 kg/m^2^) were recruited from the Max Planck Institute’s internal participants database and advertisements placed at university facilities or public spaces. All participants were non-smokers and reported no history of clinical drug or alcohol abuse or neurological disorder, and none had a first-degree relative history of psychiatric disorders. None showed moderate or severe depressive symptoms assessed with the Beck Depression Inventory, indicated by total scores < 19^45^. We decided to only include female participants since previous studies reported larger behavioral effects of APTD in women compared to men^39,46^, an effect potentially explained by higher striatal dopamine synthesis capacity in women^33^.The study was carried out in accordance with the Declaration of Helsinki and was approved by the Medical Faculty Ethics Committee of the University of Leipzig. All participants gave written informed consent before taking part in the study.

### 2.2. Study Design

Participants were first invited to the institute for a screening to check for inclusion and exclusion criteria. We used the Dietary Fat and free Sugar Questionnaire (DFS)^43,44^ to group our participants into two groups of high and low consumers of saturated fat and refined sugar (HFS vs LFS group). The DFS consists of 24 questions asking how often participants consumed a certain food item on average over the last twelve months (5 answer options; from ‘1 per month or less’ to ‘5 times or more per week’). Two additional questions ask for the frequency of food consumed outside the home averaged over the last twelve months and the number of spoons of sugar added to food and beverages in the last week. The minimum score possible is 26 (low intake of HFS) and the maximum score is 130 (high intake of HFS). DFS scores were shown to correlate with the percentage energy from saturated fat and free sugar and high intra-class correlations indicate good test-retest reliability^43,44^. Based on the total DFS score participants were assigned to the LFS group (total score < 54) or the HFS group (total score > 61); participants with DFS scores ranging from 54 to 61 were excluded from the study. Additionally, baseline fasted blood measurements were taken and participants completed the Viennese Matrices Test 2 (WMT-2) to assess intelligence^47^. They further answered self-reported questionnaires on eating behavior and personality.

If participants fulfilled all inclusion and none of the exclusion criteria, they underwent two test days with a minimum of 7 days between sessions (mean: 11.97 days, maximum: 36 days; **Fig. 1**). A within-subject, double-blind crossover design was used to test participants under a dopamine depletion condition (DEP) and a balanced dopamine condition (BAL). Test sessions were scheduled either at 8 am or 10 am, the two sessions always started at the same time for each participant. Before ingestion of the amino acid drink and at the end of the test session participants rated their well-being with digital visual analogue scales (VAS) asking for sadness, anxiety, mood, nausea, appetite, hunger, satiety, fullness and urge to move. To monitor success of the APTD intervention, blood samples were drawn before ingestion of the drink and ~ 4 h post ingestion, prior to behavioral testing. To assess working memory capacity, which is considered to be a proxy for dopamine tone^15^, the verbal forward and backward digit span task^48^ was administered in a soundproof room immediately before behavioral testing. Behavioral testing was conducted 4.5-5 h post ingestion (mean: 4 h 49 min, maximum: 5 h 38 min). During the period between ingestion and behavioral testing participants read, watched a movie or worked quietly. Two hours after ingestion participants were provided with a low protein snack, consisting of fruits (apple, banana and grapes) and vegetables (cucumber, carrots and red pepper).

**Fig. 1.**
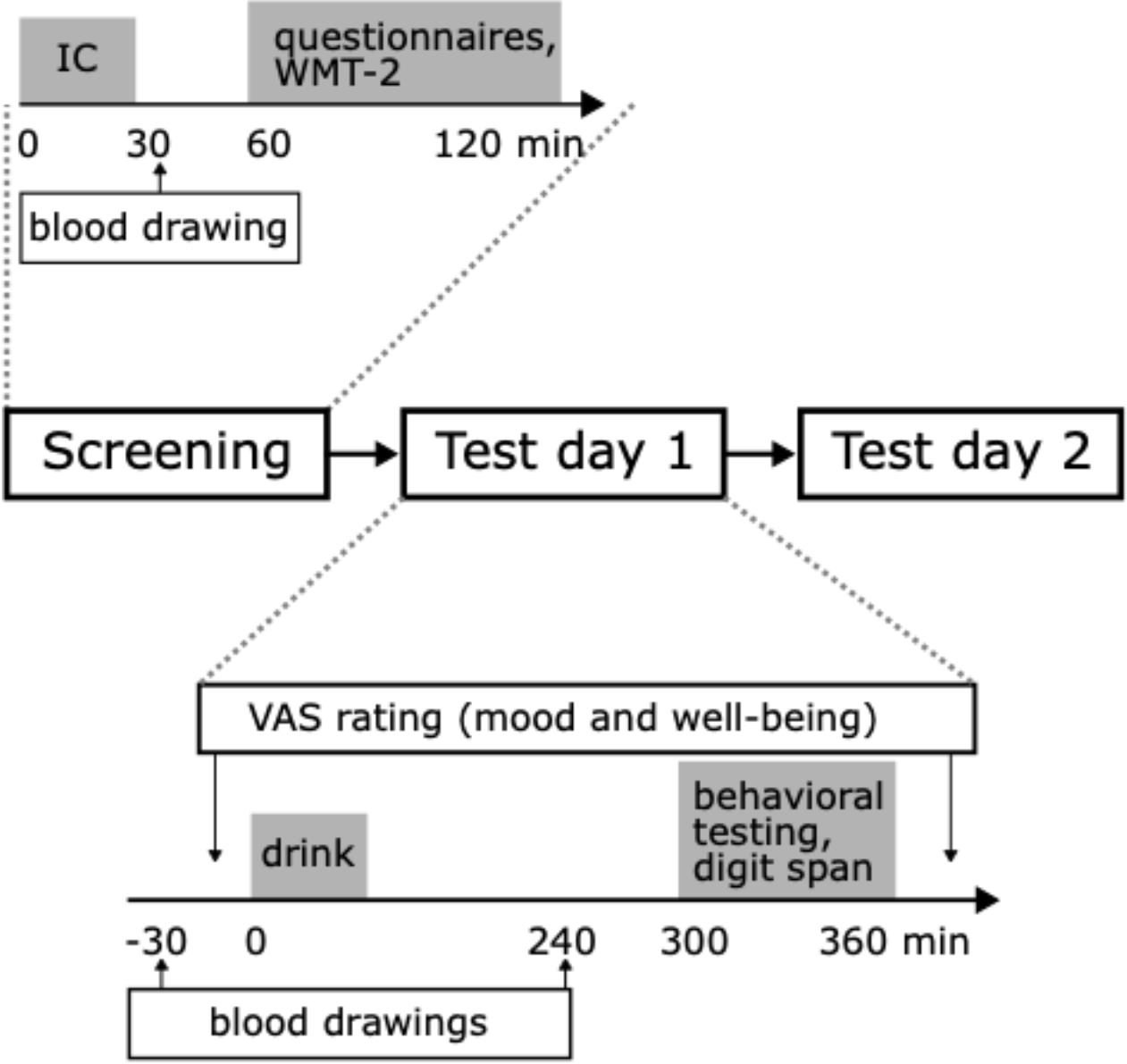
Overview of measurements and timings on screening and test day. Screening: participants gave informed consent (IC) before blood was drawn to measure pDAP at baseline. Afterwards they completed the DFS to assess study eligibility together with questionnaires for eating behavior and personality traits. Test days: participants completed two test days with different intervention drink. Behavioral testing and the digit span task were conducted ~5 h after ingestion of the intervention drink. Blood was drawn prior to ingestion of the drink and prior behavioral testing to measure pDAP pre and post intervention. Visual analog scales (VAS) were administered prior to ingestion of the drink and at the end of the test session.

### 2.3. Behavioral testing

Participants performed the Probabilistic Selection Task (PST) and the operation span task (OSPAN) as indirect measures of dopamine function on both test days. The order of PST and OSPAN was different on each test day for individual participants and randomized and counterbalanced across participants within groups. PST and OSPAN were programmed and performed using the software *Presentation 16.5* (Neurobehavioral Systems, Inc., Berkeley, CA, USA).

#### 2.3.1. Operation span task (OSPAN)

Working memory performance was examined with a modified version of the automated OSPAN task^42,49^. During the OSPAN task participants had to mentally solve a presented mathematical problem (e.g. (4*2)-7) and then indicate with a mouse click, if the presented answer is the correct answer to that problem. The time limit for answering was the average time participants needed to answer the given solutions to mathematical problems in the preceding training phase plus 2.5 standard deviations. Subsequently a target letter was presented on the screen, which participants were instructed to remember. After three to seven items (with the number of items per trial varying randomly to prevent participants from anticipating the number of items to be remembered) participants were asked to recall the items by choosing letters from a 3 x 4 matrix containing 12 letters by clicking them in the presented sequence with the mouse. Each length of items was presented three times, adding up to a total of 75 math problems and letters presented. Working memory performance was calculated using the MIS scoring method, a measure that accounts for performance on the distractor task^50^. In short, the MIS main score (referred to as *MIS score*) consists of the working memory related components ‘number of remembered items’ (I) – short-term memory – and ‘longest contiguous sequence remembered’ (S) – relative object placement – and adjusts for performance on the mathematical distractor task (M) on each trial. The MIS score for each trial was calculated using the following formula:

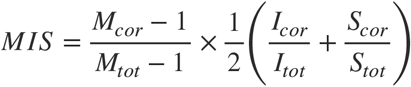

The left side of the multiplication accounts for performance on all mathematical problems (M) except the first one presented, by calculating the ratio of the number of correctly answered problems minus one to the total of mathematical problems minus 1 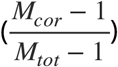. The right side of the multiplication regards short-term memory (I) and relative object placement (S) and is calculated by the ratio of number of correctly recalled items to the total number of items 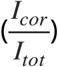 plus the ratio of the longest contiguous sequence re-called to the total length of the presented sequence 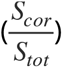. This part of the score is divided by two to weight the distractor and the working memory part of the score equally. The total *MIS score* for each participant is the sum of all scores per trial; the maximum *MIS score* possible is 15. The MIS scoring method allows to calculate a subscore only for the working memory components of the OSPAN without the distractor task by only calculating the right side of the multiplication shown above (referred to as *IS subscore*). The total *IS subscore* for each participant is the sum of all scores per trial; the maximum *IS subscore* possible is 15. The complete task with training and test phase takes around 30 minutes to finish.

#### 2.3.2. Probabilistic selection task (PST)

The PST consisted of a training and a test phase. During the training phase participants viewed three different pairs of stimuli and had to choose one of the stimuli within each pair. Stimuli consisted of six Japanese Hiragana letters (referred to as A-F), always paired as AB, CD and EF. The probabilities for positive feedback for each stimulus were predetermined (A: 80%, B: 20%, C: 70%, D: 30%, E: 60%, F: 40%). Positive feedback was signaled by the word “ correct” in green, negative feedback was signaled by the word “ incorrect” in red, both displayed centrally on the screen. Participants had 4000 ms to choose a stimulus via left or right mouse click. Failure to respond within that time led to feedback encouraging the participant to respond faster on the next trial. The stimulus pairs were presented repeatedly in random order, each pair 20 times in a block of 60 trials. After each block, learning performance was checked and if a predefined criterion was met (minimum 65% A in AB pair, 60% C in CD pair and 50% E in EF pair), participants advanced to the test phase. If the criterion was not met, participants continued with the next training block, with a maximum of 10 blocks. Participants that did not meet the criterion after 10 training blocks did not advance to the test phase and were excluded from the analysis for this task. During the test phase the six stimuli were presented in novel pairs and participants were instructed to choose the stimulus that was more likely to have been associated with positive feedback before, but this time no feedback was provided. All 15 possible combinations of stimuli were presented each four times, adding to a total of 60 trials in the test phase. Performance measures from the test phase were learning from positive feedback (“approach”; choosing A over all other stimuli and C over E and F) and learning from negative feedback (“avoid”; avoiding B in all pairs and D in when paired with E or F). Participants that failed to choose A over B in the AB pair two times or more were excluded from the analyses, since it was assumed that those participants failed to remember the reward associations of the stimulus pair that was the easiest to discriminate and were thus unable to perform the task properly. The complete task with training and test phase takes around 30 minutes to finish.

#### 2.4. Acute phenylalanine/tyrosine depletion (APTD)

To first deplete pDAP levels, participants followed a diet low in protein (< 20 g protein) on the day prior to the test sessions (guidelines provided by a nutritionist) and fasted overnight, the latest from 10 pm onwards. Drinking water was encouraged and drinking black coffee and tea (without sugar or milk) was allowed in accustomed amounts. On the BAL test days,pDAP levels were repleted by means of ingestion of an amino acid drink containing Leucine, Isoleucine, Lysine, Methionine, Valine, Threonine, Tryptophan, Tyrosine and Phenylalanine. In the DEP condition, a mixture of all aforementioned amino acids except from dopamine’s precursors phenylalanine and tyrosine was administered. The composition of amino acid mixtures was based on the formula by Mc-Tavish^51^ and adapted for three different weight classes to reach ideal dopamine depletion effects with lowest side effects; based on the formula by Frank and colleagues^52^. The three weight classes ranged from 50-67 kg, 68-83 kg, and higher than 84 kg (maximum weight 146.5 kg) and differed in total amino acid quantity but not their ratio. The amino acid drinks were mixed with lemonade (Fanta or Sprite, Coca-Cola European Partners plc, Uxbridge, United Kingdom) to cover the bitter taste and an anti-foaming agent (Espumisan, BERLIN-CHEMIE AG, Germany) for better tolerance. Successful intervention was defined as a positive difference in phenylalanine and tyrosine between post and pre intervention under the balanced condition (PheTyr_post_ – PheTyr_pre_> 0) and a negative difference between post and pre intervention under the depleted condition (Phe-Tyr_post_ – PheTyr_pre_< 0).

#### 2.5. Self-reported questionnaires

All participants completed the BDI and DFS for inclusion, and personality and eating behavior questionnaires to characterize the two diet groups on the screening day. Feeling of hunger, dietary restraint and disinhibition were assessed using the Three Factor Eating questionnaire (TFEQ)^53,54^. Personality measures encompassed the personality traits openness, conscientiousness, extraversion, agreeableness and neuroticism (NEO-FFI)^55^, behavioral inhibition and approach system (BIS/BAS)^56^ and impulsivity (UPPS)^57^. All questionnaires except for the BDI, which was administered on paper, were administered online using the statistical survey web app LimeSurvey (LimeSurvey GmbH, Hamburg, Germany).

#### 2.6. Blood measures

Blood samples for the analyses of amino acids were drawn at the screening, prior to ingestion of the drink and prior to behavioral testing on both test days. Blood samples for the analyses of metabolic parameters (cholesterol, triglycerides, glucose, glycated hemoglobin (HbA1c), insulin and leptin) were drawn at the screening and prior to ingestion of the drink on both test days; insulin resistance was calculated using the homeostatic model assessment (HOMA-IR). Whole blood samples were drawn using EDTA monovettes (2.7 ml EDTA S-monovette, SARSTEDT AG & Co. KG, Nümbrecht, Germany) and kept for 15 minutes at room temperature in upright position before being stored at −80 °C. Blood serum was drawn using monovettes with clot activator (9 ml S-monovette, SARSTEDT AG & Co. KG, Nümbrecht, Germany), kept for 30 minutes at room temperature in upright position, centrifuged for 10 minutes at 15 °C with 3500 rpm, and the supernatant stored at −80 °C. Metabolic parameters were analyzed with the COBAS 8000 system (Roche Diagnostics GmbH, Mannheim, Germany). The analysis of amino acids was performed as published previously^58,59^. In brief, for protein depletion 10 μl serum was diluted with methanol containing isotope labelled standards. After centrifugation and derivatization analysis was performed via MS/MS on an API 4500 tandem mass spectrometer (Applied Biosystems, Germany).

#### 2.7. Study samples

65 participants completed the screening day (i.e. had not to be excluded based on health issues, smoking or drug abuse), including blood drawing and self-reported measures of eating behavior and personality, and began the test days with dietary intervention. During the course of the study 16 participants dropped out voluntarily (**Fig. 2**), further three participants with an estimated IQ lower than 85 and four participants that had to vomit after ingestion of the amino acid drink on one of the test days were excluded from the analyses. Finally, 11 participants for whom the intervention was unsuccessful had to be excluded from the analyses. Statistical outliers for BMI, based on 2.2 interquartile range, were included in the analyses to ensure proficient sample size. Thus the study sample consisted of 31 subjects, 17 in the LFS group and 14 in the HFS group (**Table 1**). For analyses of the OSPAN task, one subject in the LFS group had to be excluded due to poor overall performance (statistical outlier based on 2.2 interquartile range criterion), resulting in a sample of n_LFS_ = 16 and n_HFS_ = 14 participants. During the PST, eight participants did not reach the chance criterion in the test phase, which resulted in a sample of n_LFS_ = 12, n_HFS_ = 11 participants available for analyses of this task.

**Fig. 2.**
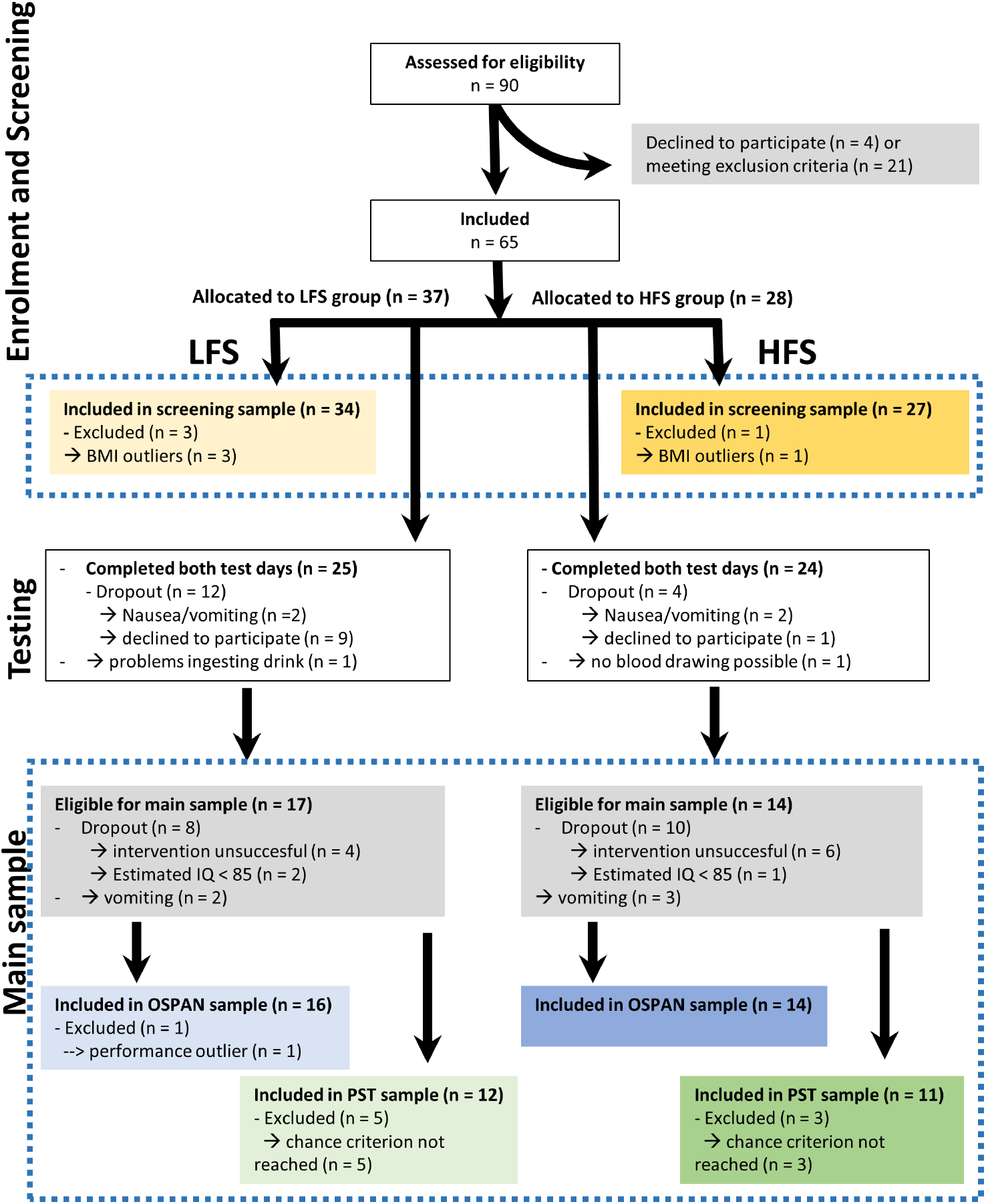
Flowchart of the study protocol. Enrolled participants were screened for eligibility based on health and diet. Included participants completed two test days varying in intervention drink. Participants with unsuccessful intervention or who vomited/felt nauseous during testing were excluded from the analyses. Task specific criteria were used to define samples for task analyses. Dashed blue frames indicate samples that were used for statistical analyses. LFS: low fat sugar; HFS high fat sugar.

**Table 1.** Group demographics and metabolic measurements of the study sample

|  | LFS<br>(n=17) |  |  | HFS<br>(n=14) |  |  | p-value |
| --- | --- | --- | --- | --- | --- | --- | --- |
|  | *M* | *(SD)* | *range* | *M* | *(SD)* | *range* |  |
| Age [years] | 24.7 | (4.3) | 19-32 | 25.8 | (4.0) | 21-33 | 0.452<sup>‡</sup> |
| BMI (kg/m<sup>2</sup>) | 26.3 | (8.5) | 20.1-50.1 | 23.1 | (4.3) | 18.5-35.9 | 0.173<sup>§</sup> |
| Non-verbal IQ<sup>†</sup> | 104.3 | (9.8) | 87-118 | 100.1 | (8.7) | 87-113 | 0.226<sup>‡</sup> |
| Cholesterol [mmol/l] | 4.5 | (0.8) | 3.2-6.4 | 4.6 | (1.0) | 3.1-6.7 | 0.552<sup>¶</sup> |
| Triglycerides [mmol/l] | 1.2 | (0.6) | 0.5-3.2 | 0.9 | (0.4) | 0.2-1.6 | 0.823<sup>§</sup> |
| HbA1c % | 5.0 | (0.2) | 4.6-5.4 | 4.9 | (0.4) | 4.4-5.5 | 0.556<sup>¶</sup> |
| Glucose [mmol/l] | 4.6 | (0.5) | 3.5-5.5 | 4.6 | (0.5) | 4.1-5.6 | 0.662<sup>¶</sup> |
| Insulin [pmol/l] | 57.3 | (41.5) | 17.1-179.2 | 45.8 | (33.5) | 13.4-119.1 | 0.765<sup>§</sup> |
| HOMA-IR | 2.0 | (1.4) | 0.5-5.8 | 1.6 | (1.3) | 0.4-4.8 | 0.707<sup>§</sup> |
| Leptin [ng/ml] | 17.7 | (20.8) | 0.2-90.2 | 15.3 | (16.8) | 4.6-70.2 | 0.568<sup>§</sup> |

Abbreviations: n = number of participants; M = mean ; SD = standard deviation; BMI = body mass index in kg/m<sup>2</sup>; † Non-verbal IQ calculated based on the Viennese Matrices Test 2, ‡ unpaired two-sample *t*-test, § Wilcoxon signed-rank test, ¶ ANCOVA

Because a secondary aim of this study was to characterize the two dietary groups with respect to the dopaminergic system, but also metabolic parameters, eating behavior and personality, we extended the analyses of measurements obtained at screening day to all participants that completed the screening day. In this screening sample we excluded four BMI outliers based on 2.2 interquartile range criterion for the analyses of physiological (pDAP and metabolic parameters) and questionnaire data to minimize the effect of body weight on those measures. As a result, the analyzed screening sample consisted of 61 participants, 27 in the HFS group and 34 in the LFS group.

#### 2.8. Statistical Analyses

Statistical analyses were performed in R v3.4.3^60^ within RStudio^61^, using the packages *car, stats, pastecs, nlme*, and *psych*. Unpaired two-sample *t*-test or Wilcoxon signed-rank test, when appropriate, were used to analyze the group demographics age, BMI and IQ, and questionnaire data. All data that was measured multiple times (cognitive task performance, digit span, pDAP,VAS) was analyzed with a linear mixed model, simple effects analyses were used to test the direction of significant interactions. All tests involving cognition (OSPAN, PST, digit span) were controlled for age, IQ, and BMI by adding those as covariates of no interest in the linear mixed model. Performance of the OSPAN task (*MIS score* and *IS subscore*) was analyzed using a three-way interaction model with the factors ‘diet group’ and ‘intervention’ and the covariate of interest ‘pDAP at screening’ (as proxy for the dopaminergic system). Predictor variables for the analyses of the training phase of the PST were ‘diet group’ and ‘intervention’, predictor variables for the analyses of the test phase were ‘diet group’, ‘intervention’ and ‘test condition’ (approach vs. avoid). Reaction times of the OSPAN and PST were analyzed in *diet group x intervention* models, correcting only for age. Working memory capacity (digit span forward, backward, total) was analyzed with a *diet group x intervention* model. Metabolic blood parameters from screening day and pDAP at all time points were controlled for age and BMI. Metabolic blood parameters were tested using analysis of covariance (ANCOVA) or by regressing out the variance explained by the covariates and performing group comparisons on the residuals when data did not meet the assumptions for ANCOVA (Wilcoxon signed-rank test). To test how pDAP changed between time points on test days, we ran a model with ‘diet group’, ‘time point’, and ‘intervention’ as independent variables. We further tested if pDAP was different between diet groups and interventions at baseline (screening and pre ingestion of the drink at test days) with a *diet group x intervention* model. The contrast ‘screening’ vs. ‘test days’ within the factor ‘intervention’ of that model was used to test the effect of the 24 h low-protein diet the day prior testing. Diet group differences in pDAP at screening and the baseline of test days (mean of both test days) were tested separately using ANCOVA with age and BMI as covariates or by regressing out the variance explained by the covariates and performing group comparisons on the residuals when data did not meet the assumptions for ANCOVA (Wilcoxon signed-rank test). Independent variables for the analysis of VAS were diet group (LFS vs. HFS), time point at test day (pre vs. post) and intervention (BAL vs. DEP). In all models, lower level interactions and main effects were only regarded when the higher order interactions were not significant. The significance level alpha was 0.05 unless stated differently when corrected for multiple comparison. All effect sizes are reported as Pearson r.

Absolute values of amino acid levels were z-transformed, due to batch differences in the ranges of values of the analyzed samples in the lab (samples were send to the lab at two different time points). For comparing precursor availability at the different baseline measurements (screening and test days) and pre and post intervention, the absolute values were z-transformed using the mean and standard deviation of all 5 measurements in each batch. The values of the screening sample were z-transformed using the mean and standard deviation of each batch at screening day.

## 3. Results

This study was designed to investigate the differential effects of a dietary dopamine depletion depending on low vs. high self-reported intake of HFS on performance on a working memory and reinforcement learning tasks – as indirect measures of dopamine function. First we checked whether the intervention was successful in our sample by comparing pDAP before and after the intervention. BAL and DEP intervention changed pDAP in opposite directions (*χ^2^*(1) = 207.78, *p*< 0.001, r = 0.920) independent of diet group (χ^2^(1) = 0.361, *p* = 0.548, r = 0.057); ingestion of the BAL drink increased pDAP (*χ*^2^(1) = 161.61, *p* < 0.001) and ingestion of the DEP drink reduced pDAP (*χ^2^*(1) = 454.32, *p* < 0.001; **Fig. 3**). Consequently pDAP was significantly lower after ingestion of the DEP compared to the BAL drink (*χ^2^*(1) = 1088.65, *p* < 0.001).

**Fig. 3.**
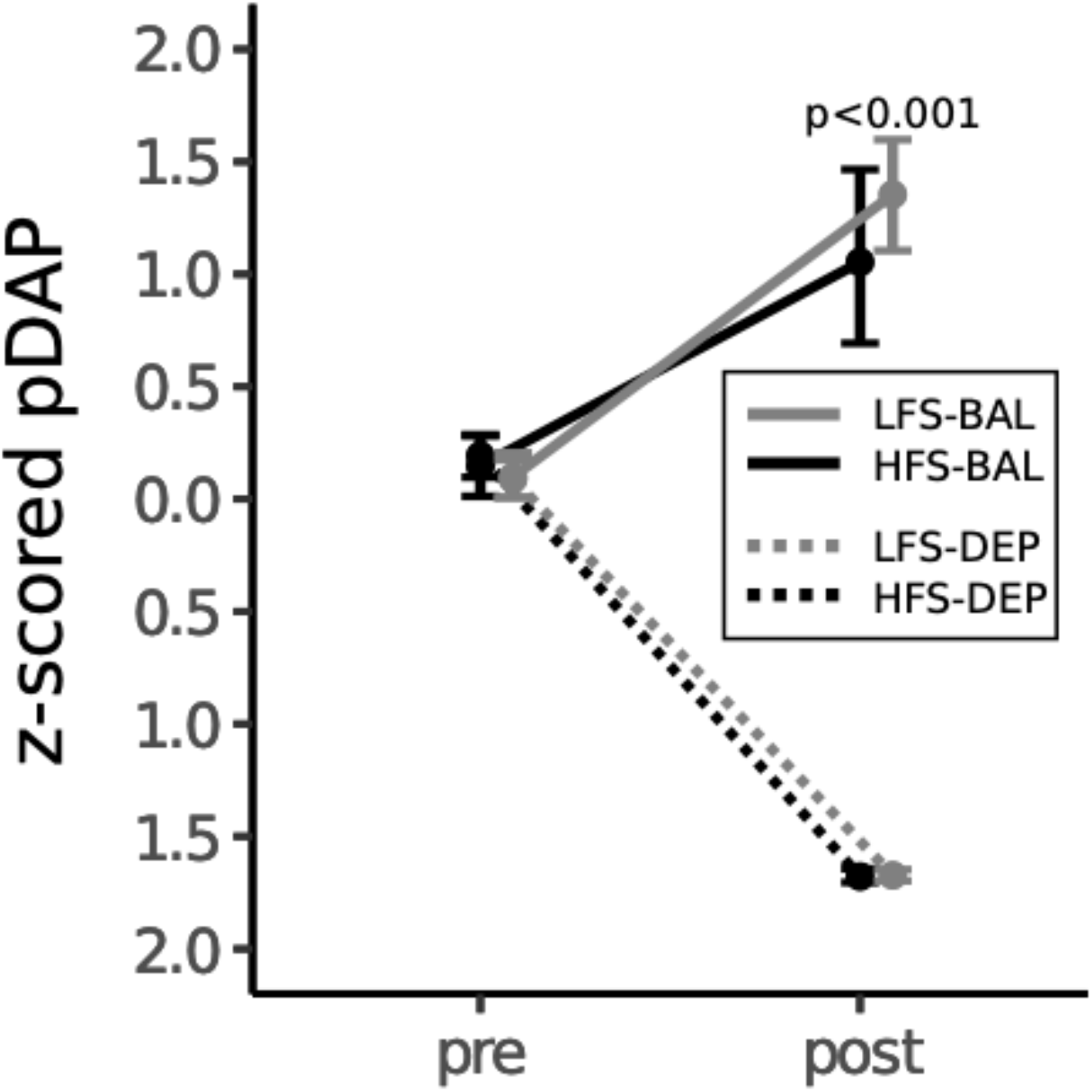
pDAP pre and post ingestion of the amino acid drinks. ingestion of the BAL drink increased pDAP (*χ^2^*(1) = 161.61, *p* < 0.001) and ingestion of the DEP drink reduced pDAP (*χ^2^*(1) = 454.32, *p* < 0.001. pDAP was significantly lower after ingestion of the DEP compared to the BAL drink (*χ^2^*(1) = 1088.65, *p* < 0.001). nLFS = 17, nHFS = 14.

### 3.1. Effects of dopamine manipulation on cognitive performance

#### 3.1.1. Working memory

On each test day we measured working memory capacity with the forward and backward digit span task to account for possible effects of the intervention on diet groups and group differences in working memory capacity that might explain different performance on the OSPAN task. The intervention did not affect working memory capacity (forward, backward and total) in the two diet groups differently, and it did not differ between diet groups and intervention (all *p*> 0.85).

Working memory performance was tested with the OSPAN task and scored using the method proposed by Lammert and colleagues^50^. According to this method, the *MIS score* of the OSPAN task additionally regards performance on the mathematical distractor task, whereas the *IS subscore* is supposed to reflect the working memory component of this task only. Therefore, we focus on the *IS subscore* as measure for working memory performance. First, we analyzed the effects of APTD on the complete *MIS score*. Overall both diet groups did not differ in *MIS score* (*χ^2^*(1) = 2.764, *p* = 0.096, r = 0.117), but were similarly impaired after ingestion of the DEP drink at trend level (*χ^2^*(1) = 3.56, *p* = 0.059, r = 0.467; **Fig. 4A**). Looking at the working memory-specific *IS subscore* the intervention had differential effects on working memory performance on the two diet groups (*χ^2^*(1) = 5.19, *p* = 0.023, r = 0.009; **Fig. 4B**). Simple effects analyses showed that after ingestion of the DEP drink the LFS group performed worse than after ingestion of the BAL drink (*χ^2^*(1) = 9.73, *p* = 0.002) and worse than the HFS group after ingestion of the DEP drink (*χ^2^*(1) = 6.83, *p* = 0.009). Furthermore, the intervention affected participants’ working memory performance depending on their pDAP at screening day independent of diet group: participants with lower pDAP at baseline performed better under the BAL condition and got worse under the DEP condition, whereas participants with higher pDAP performed slightly better under the DEP than under the BAL condition(*χ^2^*(1) = 8.22, *p* = 0.013, r = 0.414; **Fig. 4C**). Because dopamine is also involved in motor control and movements we additionally analyzed reaction times of the cognitive tasks. Reaction times for solving the mathematical problem, evaluating the presented solution, and recalling the letter sequence of the OSPAN did not differ significantly between diet groups or intervention (all *p*> 0.85).

**Fig. 4.**
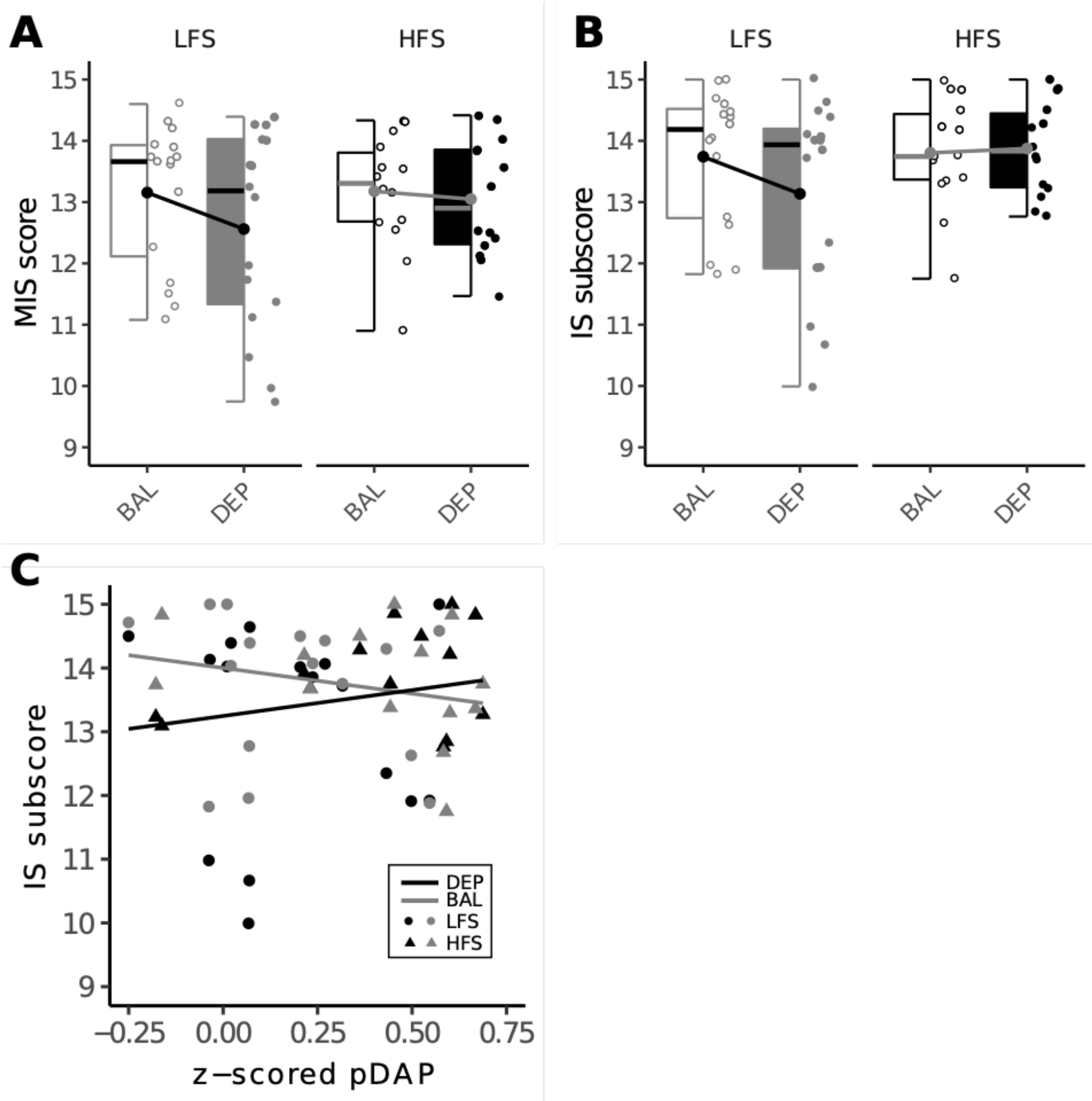
Effects of APTD on working memory performance measured with the OSPAN. **A)** MIS score for the two dietary groups underbalanced (BAL) and depleted (DEP) condition. DEP trend significantly decreased working memory performance in both groups (*χ^2^*(1) = 3.56, *p* = 0.059, r = 0.323). **B)** MIS subscore that stronger represents the working memory component of the OSPAN. The DEP condition impaired working memory performance in the LFS group but did not affect performance of the HFS group (*diet group x interventionχ^2^*(1) = 5.19, *p* = 0.023, r = 0.210). **C)** Relationship of peripheral dopamine precursor availability (pDAP) at screening and working memory measured with the MIS subscore. Grey symbols and regression line represent BAL condition, black regression line and symbols represent DEP condition; dots represent LFS group, triangles represent HFS group. Participants with lower pDAP performed better under the BAL than the DEP condition, whereas participants with higher pDAP were merely affected by the depletion (*pDAP x intervention*χ^2^(1) = 8.22, p = 0.004, r = 0.499). nLFS = 16, n HFS = 14.

#### 3.1.2. Reinforcement learning

Reinforcement learning was tested with the PST which consists of a training and a test phase^14^. Learning of reward associations during the training phase did not differ between diet groups (*χ^2^*(1) = 0.027, p = 0.869, r = 0.036), but participants in both groups required more training blocks to reach the learning criterion after ingestion of the DEP drink ((*χ^2^*(1) = 3.63, *p* = 0.057, r = 0.499; **Fig. 5A**). The test phase of the PST tests how well participants learned to approach rewarded stimuli and avoid punished stimuli (referred to as test condition). APTD increased accuracy of choices in the test phase (*χ^2^*(1) = 3.916, *p* = 0.048, r = 0.301), independent of DFS group (*χ^2^*(1) = 0.137, *p* = 0.711, r = 0.126) and test condition (*χ^2^*(1) = 0.051, *p* = 0.822, r = 0.076, (**Fig. 5B**). To check if better performance under the depleted condition was mediated by the higher number of training blocks, we correlated the number of training blocks and accuracy in the test phase (mean of both test conditions) for the BAL and DEP condition separately. Number of training blocks and accuracy on the test phase did not correlate significantly in neither the BAL nor the DEP condition (Kendall’s *τ* DEP = 0.206, Kendall’s *τ* BAL = −0.113, *p*-values > 0.201). Reaction times did not differ significantly between diet groups or intervention conditions in the training phase (all *p*> 0.657), but the HFS group got slower and the LFS faster after ingestion of the DEP drink in the test phase (*χ^2^*(1) = 3.407, *p* = 0.065, r = 0.367).

**Fig. 5.**
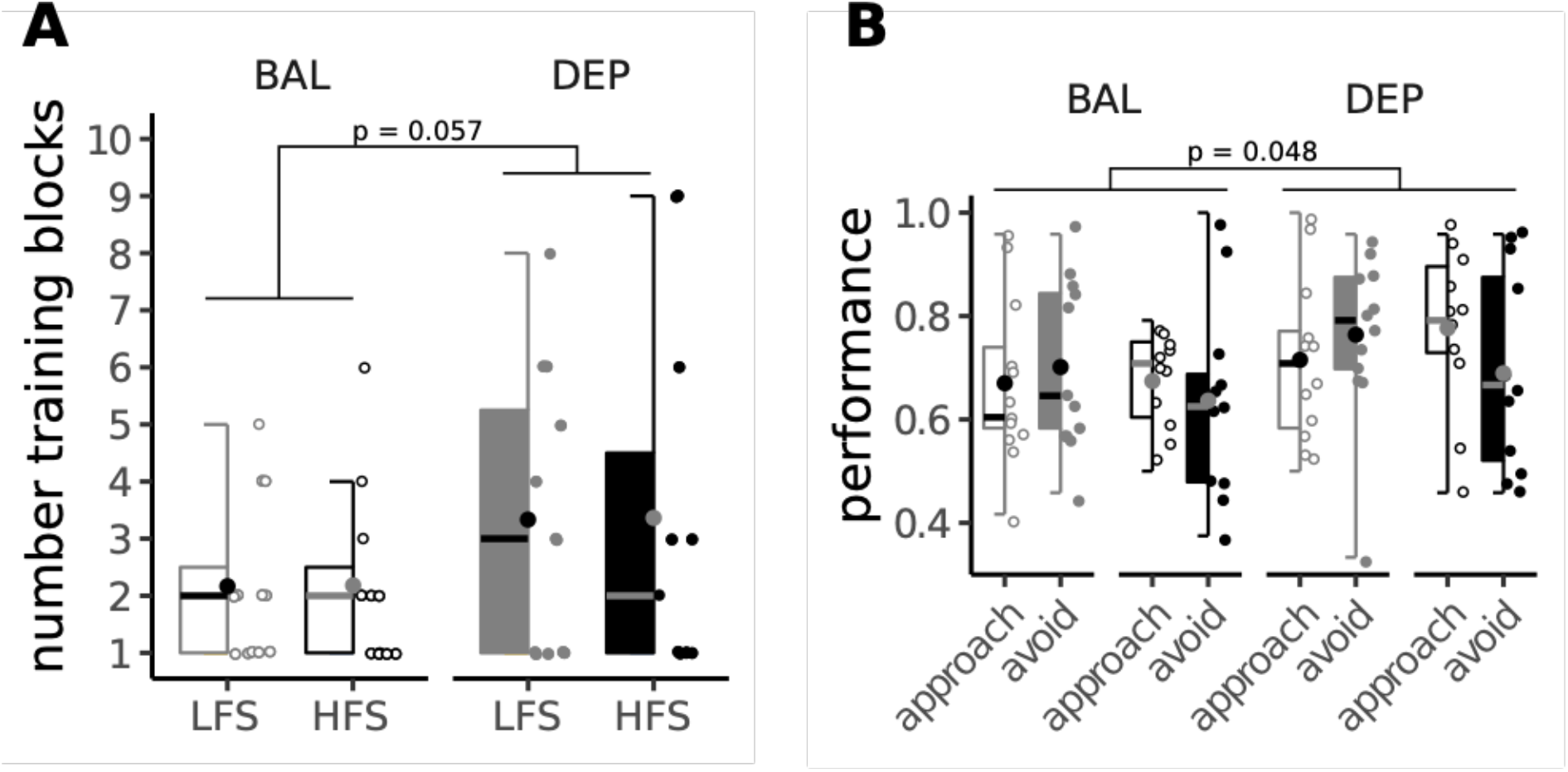
The effect of APTD on reinforcement learning. **A)** Number of training blocks necessary to advance to the test phase of the PST. Both groups needed trend significantly more training blocks under the DEP condition to reach the learning criterion and advance to the test phase (*χ^2^*(1) = 3.63, *p* = 0.057, r = 0.499. **B)** Performance in the test phase of PST. Depletion significantly increased the performance in both the approach and avoid condition in both diet groups; *χ^2^*(1) = 3.916, *p* = 0.048, r = 0.144. nLFS = 12, nHFS = 11.

#### 3.1.3 The effects of dopamine-depletion on mood and well-being

To check if dopamine depletion affects potential confounders such as mood and wellbeing differentially in the two diet groups, we analyzed scores on visual analog scales that were employed before the ingestion of the drink and at the end of each test day. Ingestion of the BAL and the DEP drink did not change any measure of mood or well-being differently in the two groups (all *p*> 0.100). Sadness and anxiety scores were increased at the end of the test day when participants had ingested the DEP drink, but decreased after ingestion of the BAL drink (sadness: *χ^2^*(1) = 4.19, *p* = 0.041, r = 0.141; anxiety: *χ^2^*(1) = 5.11, *p* = 0.024, r = 0.119). Fullness, nausea, satiety and the urge to move were increased after ingestion of the amino acid drink on both test days (all *p* < 0.006). We ran the same statistical models with the order of test days (day 1 or day2) instead of intervention type as factor to check whether experiences from the first test day affected participants’ well-being and mood irrespective of the intervention. Only sadness decreased over the course of the first test day but increased during the second test day; (*χ^2^*(1) = 6.29, *p* = 0.012, r = 0.170).

### 3.2 Peripheral dopamine precursor availability

We assessed group differences in pDAP at baseline (screening, prior ingestion of amino acid drinks on test days) as a proxy for the status of the central dopamine system in the two dietary groups, since constantly higher levels of dopamine could induce the alterations of dopaminergic transmission observed in rodents after HFS intervention^22–29^. pDAP was significantly higher in the HFS relative to the LFS group when tested over all three baseline measurements (*χ^2^*(1) = 5.50, *p* = 0.019, r = 0.484; **Fig. 6**). Because participants’ diet prior to test days (low protein diet) differed compared to the diet prior to the screening day (regular diet) and this low protein diet lowered pDAP compared to the screening day in both groups (*t*(58) = −4.235, *p* < 0.001, r = 0.484), we also compared pDAP between both diet groups separately at screening and test sessions (averaged across BAL and DEP day). This analysis revealed that the HFS group had higher pDAP at the screening session after having followed a regular diet (*W* = 69, *p* = 0.048, r = −0.355), and only slightly higher on the test days after having followed a one-day diet low in protein (*F*(1,28) = 3.29, *p* = 0.081, r = 0.324). This group difference in pDAP supports our finding that APTD impairs working memory performance in the LFS group as well as in participants with lower pDAP at baseline.

**Fig. 6.**
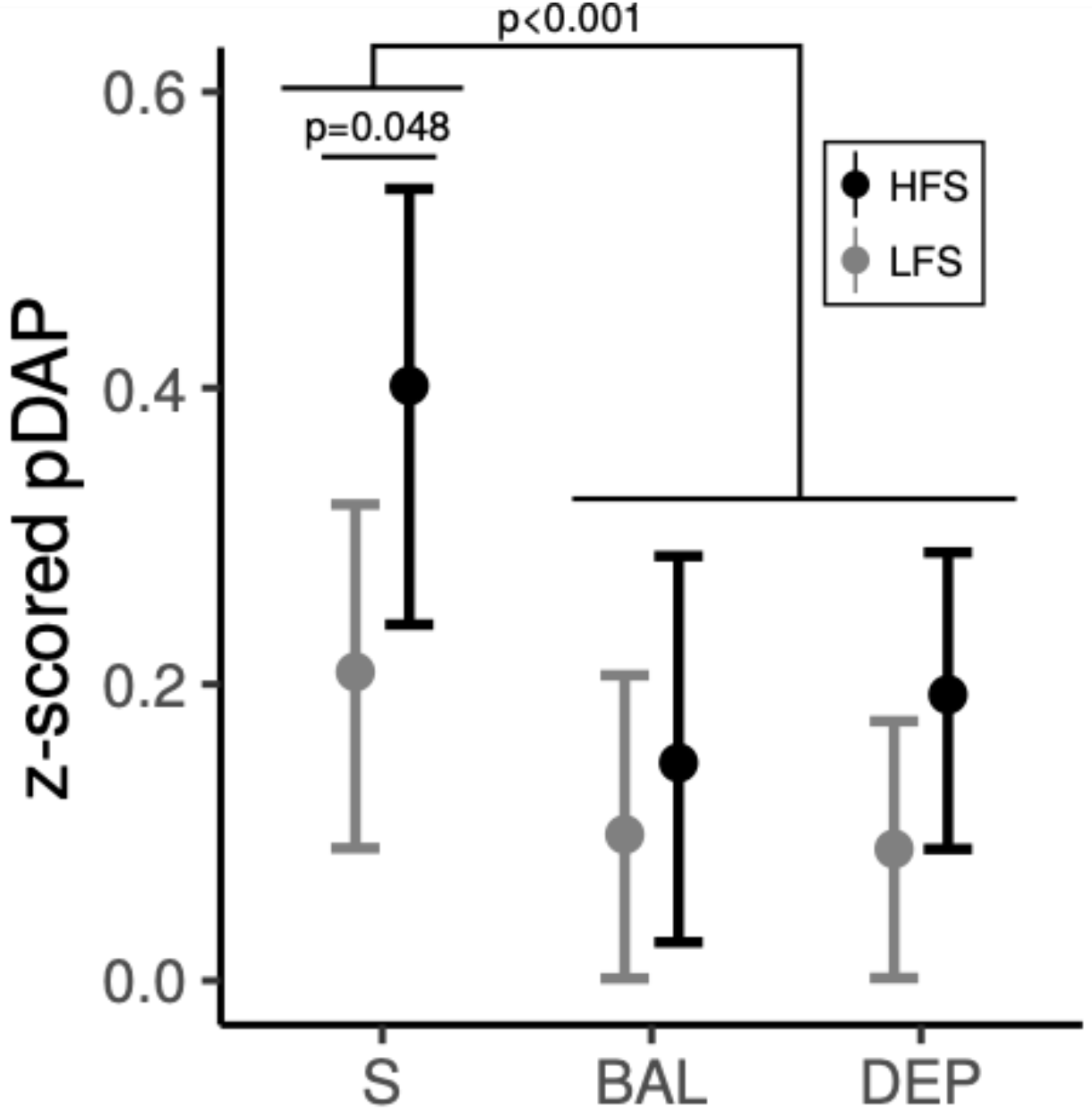
pDAP measured at the screening day (S) and pre ingestion of the intervention drinks (BAL and DEP) The low protein diet the day before test days reduced pDAP compared to the screening day with normal diet the day before (*t*(58) = −4.235, *p* < .0001, r = 0.484). The HFS group had significantly higher pDAP at the screening; *W* = 69, *p* = 0.048, r = −0.355) and trend elevated levels at baseline of test days (*F* (1,28) = 3.29, *p* = 0.081, r = 0.324).

Because pDAP and working memory capacity measured with the digit span task are both considered as proxies for central dopamine levels, we tested the correlation between these two measures. pDAP at baseline did not correlate with mean (BAL and DEP) forward, backward or total digit span (all *p*> 0.894), and pDAP prior to ingestion of the intervention drink did not correlate with forward, backward or total digit span on that test day (BAL and DEP: all *p*> 0.346).

### 3.3 Self-reported eating behavior and personality traits

Because the preference for HFS might be influenced by general differences in eating behavior, we investigated potential group differences on the three factor eating questionnaire. The HFS group showed significantly lower restrained eating (*W* = 66, *p* = 0.035, r = – 0.379), higher disinhibition (*W* =174, *p* = 0.029, r = −0.392), and higher hunger feeling (*W* = 179, *p* = 0.017 r = −0427).

Furthermore we examined if the frequency of consumption of HFS is associated with personality traits. The HFS group scored higher at trend level on the neuroticism subscale of the NEO-FFI (*t*(29) = 1.925, *p* = 0.064, r = 0.337), and on the urgency subscale of the UPPS (*t*(29) = 1.91, *p* = 0.066, r = 0.334).

### 3.4 Metabolic blood parameters

We analyzed parameters of the fat and sugar metabolism and eating related hormones to check if the dietary preference of the groups is reflected in physiological measurements. The two dietary groups did not differ in any parameter of fat (cholesterol and triglycerides) or sugar metabolism (glucose and HbA1c), as well as leptin, insulin and insulin resistance (all *p*> 0.556; **Table 1**).

### 3.5 Further characterization of diet groups (screening sample)

Because an aim of this study was to characterize the two dietary groups with respect to the dopaminergic system, but also metabolic parameters, eating behavior and personality, we extended the analyses of measurements obtained at screening day to all participants that completed the screening day and were not excluded based on health issues (supplementary Table S1). Due to the higher sample size we did exclude four statistical outliers for BMI for a more homogeneous sample (three in LFS and one in HFS group).

The two groups in the extended sample did not differ in BMI (*W* = 459, *p* = 1, r <0.001) and IQ (*W* = 443, *p* = 0.819, r = 0.029). However, age was significantly higher in the HFS group relative to the LFS group (*W* = 602, *p* = .038, r = 0.266). In this extended sample pDAP was only trend significantly higher in the HFS group (*F*(1, 57) = 3.019, *p* = 0.088, r = 0.224). Similar to the original sample the two groups did not differ in any of the metabolic parameters (all *p*> 0.14), except for a difference at trend level in cholesterol (*W* = 581, *p* = 0.078, r = −0.226). Group comparisons for the questionnaires assessing eating behavior and personality traits were corrected for age, because of the group difference in age in this sample. The HFS group still showed lower restraint eating (*W*=244, *p* = 0.003, r **=** −0.382), and higher hunger feeling (*W*=586, *p* = 0.037, r **=** −0.266). The higher disinhibition of the HFS relative to the LFS group was not observed in this larger sample (*W*=539, *p* = 0.165, r **=** −0.178). The higher neuroticism score of the HFS relative to the LFS group was now significant (*F*(1,57) = 5.66, *p* = 0.021, r **=** −0.300) and the score on the urgency subscale of the UPPS remained higher for the HFS relative to the LFS group at trend level (*F*(1,57) = 3.05, *p* = 0.086, r = 0.226). Furthermore the HFS group showed significantly higher agreeableness than the LFS group (*W*=599, *p* = 0.023, r **=** −0.291). The groups did not differ in any of the other measures of eating behavior and personality trait (all *p*> 0.165).

## Discussion

We aimed to collect first evidence that dietary intake of saturated fat and added sugar is associated with alterations of the dopaminergic system in humans. For this purpose, we grouped participants based on their self-reported fat and sugar intake into a low and a high consumer group (LFS vs. HFS). In a within-subjects design we investigated the diet-dependent effects of central dopamine depletion (using APTD) on dopamine-mediated cognitive performance in a working memory task (OSPAN) and a reinforcement learning task (PST). Furthermore, the groups were characterized in terms of peripheral dopamine precursor availability (pDAP; a proxy for dopamine system status), metabolic parameters, eating behavior and personality traits at baseline.

The main findings of this study are (1) different levels of pDAP at baseline and (2) the differential effect of APTD on working memory performance (the *IS subscore*) in the two diet groups. More specifically, we show that central dopamine depletion led to decreased working memory performance in the LFS group, whereas performance was unaffected in the HFS group. Work by Cools and D’Esposito^15^ suggests the existence of an inverted-u shaped relationship between action of dopamine levels and human working memory, with an optimum level of dopamine for performance. Along those lines, the observed differential effect of APTD on working memory may indeed indirectly reflect underlying group differences in dopamine transmission. Reducing central dopamine levels with APTD^35,36^ shifts both groups on the proposed inverted-u function, either further away from or closer to the optimum dopamine level, which is reflected in task performance (**Fig. 7**). Performance of the LFS group is reduced after APTD, which suggests the LFS group is located on the left-hand side of the inverted-u function. Performance of the HFS group at a similar level as the LFS group under the BAL condition and no change in performance after depletion suggests that the HFS group may have started on the right-hand side of the optimum. Thus the reduction of pDAP, that was similar in both groups, might have shifted the HFS on a less steeper part of the inverted-u, resulting in no measurable change in performance. Possible molecular explanations for the absence of an effect of the depletion in the HFS group could be that the putatively higher dopamine level of the HFS group (a) induces compensatory structural changes at dopaminergic synapses, like altered receptor or transporter expression, (b) expressed receptors become more sensitive to binding ligands^63^ or (c) the HFS group has a higher central capacity to buffer dopamine and hence withstand peripheral depletion. Due to the applied intervention that revealed possible differences in dopaminergic system status associated with HFS, we cannot make any statement about or draw inferences from dopamine dependent cognition at baseline. Successful intervention required participants in both groups to adhere to the same low-protein diet prior to the test days which differed from their normal diet. This might temporarily have rendered the groups more similar, in contrast to being tested on their normal diet. Thus it is difficult to compare our findings with the only other study so far, looking at the association of HFS with PFC-related executive function in humans, by Francis and Stevenson^64^, who found no differences between LFS and HFS groups. The authors argued that the effects of diet on PFC may be too subtle to detect with the administered tasks. Thus we propose that further studies with more dopamine-sensitive tasks are warranted to investigate possible baseline differences in dopamine dependent cognition associated with HFS.

**Fig. 7.**
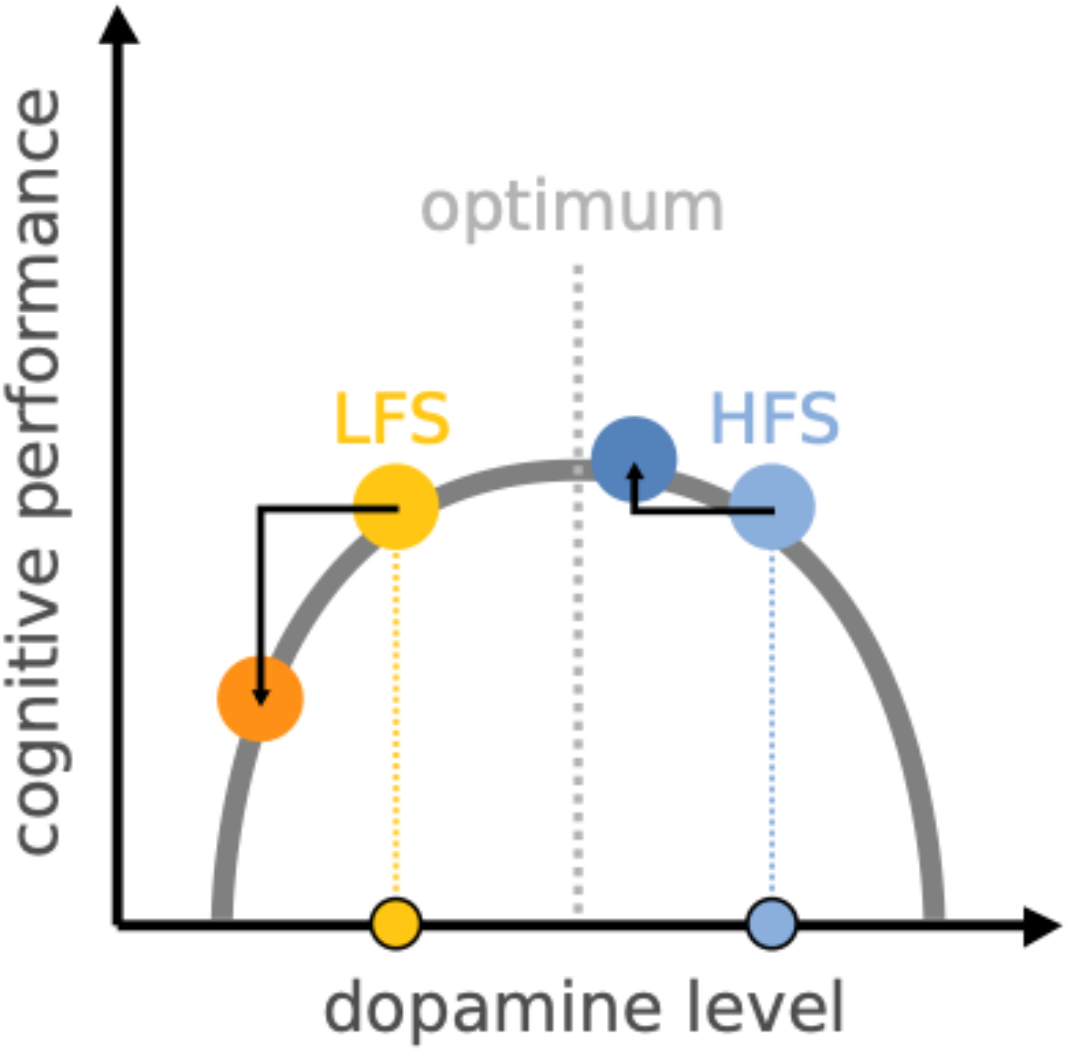
Proposed dopamine levels of the two dietary groups under BAL and DEP condition. Based on the differential effect of APTD on working memory performance (MIS subscore) we propose higher central dopamine levels in the HFS group at baseline. APTD shifts both groups on the inverted-u shaped curve of performance to lower dopamine levels. Performance in the LFS group was decreased after APTD, i.e. the group was shifted away from the optimum, which puts the LFS group on the left side of the in-verted-u curve. Performance of the HFS group was unchanged when dopamine levels were decreased, either because the inverted-u has a plateau around the optimum and this group was shifted on that plateau or the group was shifted beyond the optimum and ended up where the LFS group started.

Contrary to working memory performance, APTD affected performance on the PST equally in both groups. Independent of group, participants needed more training blocks to reach the learning criterion following APTD. In the test phase however, participants had higher accuracy after APTD. The better test performance did not relate to more training, as shown by the non-significant correlation of training blocks and test performance, suggesting that APTD has opposing effects on outcome-based learning and choice performance independent of valence. This is in contrast to other studies using reinforcement learning tasks and dopamine manipulations that either report effects on choice behavior in the test phase with unaffected initial learning^65,66^ or no mention of initial learning performance at all^14,38^. The fact that we observe no significant association of diet with performance on the PST, nor an interaction between diet and the intervention has to be interpreted with care. One important limitation of the current study is the small sample that we could analyse for this particular task, which negatively affects the power to detect diet-related differences. Furthermore, the retest reliability of the PST – together with other tasks tapping into self-regulation – has recently been called into question by a large scale literature review and empirical study by Enkavi and colleagues^67^, which is highly relevant for within-subject designs as ours. Theoretically, subtle differences in variance in task performance explained by the interaction between diet and dopamine depletion could therefore be masked by the random variance inherent to the task.

Interestingly, the observation of increased pDAP in the HFS relative to the LFS group also points to the potential existence of diet-related group differences in central dopamine transmission. Studies applying APTD showed that peripheral availability of the two dopamine precursor amino acids phenylalanine and tyrosine correlated with dopamine release in the brain^35,36^. Given the additional observation that a 24-h low protein diet prior to the test days reduced pDAP, our findings raise the hypothesis that diet-related differences in central dopamine, like they have been observed in animals after HFS intervention, may very well be the result of acute dietary effects on peripheral amino acid levels. Such short term effects have been shown for the ratio of carbohydrates to protein in a standardized breakfast^68^. It should be noted that the current study did not include a direct measure of central dopamine such as PET and can therefore not confirm our hypothesis. In addition, long-term dietary interventions are required to confirm an effect of HFS diets on amino acid level availability and dopamine-mediated cognition, preferably including PET-measurements of central dopamine levels in humans. Another possible explanation for the observed group difference in pDAP could be that the LFS and HFS group differ in the absorption or metabolism of phenylalanine and tyrosine compared to the other LNAAs and the resulting differences in central dopamine in turn could influence the preference for high-caloric food. Such a mechanism could explain why the groups still differed slightly in pDAP after they consumed a comparable diet for 24 h. This hypothesis is highly speculative though, since all LNAAs, including phenylalanine and tyrosine, are transported by the same transporter in the gut^69^ and altered amino acid metabolism can lead to severe diseases like phenylketonuria^70^. To our knowledge, only one study looked at the relationship of central dopamine transmission and food preference in humans. Wallace and colleagues combined a food rating paradigm, asking for wanting and perceived healthiness of various food items, with PET to measure striatal dopamine synthesis binding^71^. They report higher preference for perceived healthy, but not objectively healthy food items in people with lower striatal dopamine synthesis, supporting the hypothesis that endogenous dopamine is indeed related to food preference.

In our main sample as well as in the extended screening sample we found group differences in eating behavior and personality traits. In line with Francis and Stevenson^64^, who used similar dietary groups, the LFS group reported significantly higher dietary restraint, and lower hunger than the HFS group. Additionally, the HFS group in our study reported significantly higher disinhibition, though only in the smaller main sample. If eating behavior itself influences dopamine dependent cognitive performance has only been investigated in one study so far. Just recently Sadler and colleagues reported lower working memory capacity measured with the n-back task for participants with higher self-reported dietary restraint and between group differences in reward and punishment sensitivity measured with the PST^62^. To disentangle possible effects of diet and eating behavior, diet intervention studies with two groups differing in any of the TFEQ subscales are needed. Our two dietary groups also differed in the two subscales neuroticism and agreeableness of the NEO-FFI and the urgency subscale of the UPPS. Food preference and dietary style have been associated with personality traits before: in line with our finding, higher neuroticism is associated with higher preference for and consumption of sweet foods^72,73^. But it is still debatable whether personality traits influence food consumption^74^ or if more basal factors like genetic predisposition are stronger contributors^75^.

Note that the dopamine depletion effects have to be interpreted with care and await future replication. Sample size was low due to an unusual high dropout rate (**Fig. 1**) compared to other studies that administered APTD^36,38^. Our administration of the APTD intervention differed in the sense, that we mixed the amino acid drink with lemonade instead of syrup, and this syrup might have a stronger flavour to disguise the bitter taste of amino acids. Furthermore, other studies administered the unpleasant amino acids like methionine separately from the dissolved mixture^76,77^ to reduce risk of nausea. We recommend that future APTD studies follow these precautions. Furthermore generalizability of our findings is limited since we only included young healthy women in this study and we cannot make any statement about possible interaction effects of HFS and obesity, because our sample included participants from the normal weight to obese range. We also know that the genetic background influences baseline dopamine transmission parameters and cognitive function^78^, which we cannot exclude in our study. Future studies including men and women, focusing on a more narrow range of BMI and with a sample size large enough to consider genotypic variation affecting dopaminergic transmission are needed to shed further light on the association of HFS and the dopaminergic system in humans.

## 5. Conclusions

This study provides first evidence that the amount of saturated fat and refined sugars habitually consumed is associated with alterations of indirect markers for the dopaminergic system in humans. We could show that (1) the effect of a dietary dopamine depletion on working memory (but not reinforcement learning) performance, and (2) peripheral availability of dopamine precursors, a proxy for central dopamine levels^35,36^, differed between two groups who reported high relative to low intake of high fat and sugar food products, independent of body weight.

## Acknowledgements

We thank Arno Villringer and his coworkers of the Department of Neurology (MaxPlanck Institute for Human Cognitive and Brain Sciences, Leipzig, Germany) for providing lab space and additional resources. We further thank SusePrejawa for administrative support, Ramona Menger and Sylvia Stasch for preparing the intervention, and Bettina Johst and Steven Kalinke for cognitive task programming. Furthermore we thank Emmy Kaspar for participating in the preparation of the study, Lisa Ulbrich, Pauline Baßler, and Denise Linke for testing participants. Special thanks to Mathis Lammert, Linda Grasser, Lisa Leyendecker, Franziska Schwachheim for performing the blood drawings and physiological measurements. We thank Lydia Hellrung for helpful input regarding the APTD procedure. The authors have stated explicitly that there are no conflicts of interest in connection with this article. This work was funded by the Deutsche Forschungsgemeinschaft (DFG, German Research Foundation) – project number 209933838 – SFB 1052, subproject A5 (to AH, HH) and the Integrated Research and Treatment Center Adiposity-Diseases, Federal Ministry of Education and Research (BMBF), Germany FKZ: 01E01501 (to AH, LKJ, LP).

## Data Availability Statement

The data that support the findings of this study are available from the corresponding author upon reasonable request.

## Author Contributions

H.H.; Performed experiments, analyzed data and co-wrote the paper. L.K.P.; Designed and performed experiments, analyzed data and provided feedback for the paper. L.K.J.; Designed experiments, analyzed data and co-wrote the paper. S.H.; developed the lowprotein diet plan for successful intervention. U.C.; coordinated and supervised the analyses of amino acids and metabolic parameters. A.H.; supervised the research, conceived the original idea, designed experiments, helped interpreting data and provided feedback for the paper.

## Supplementary Materials

**Table S1.**
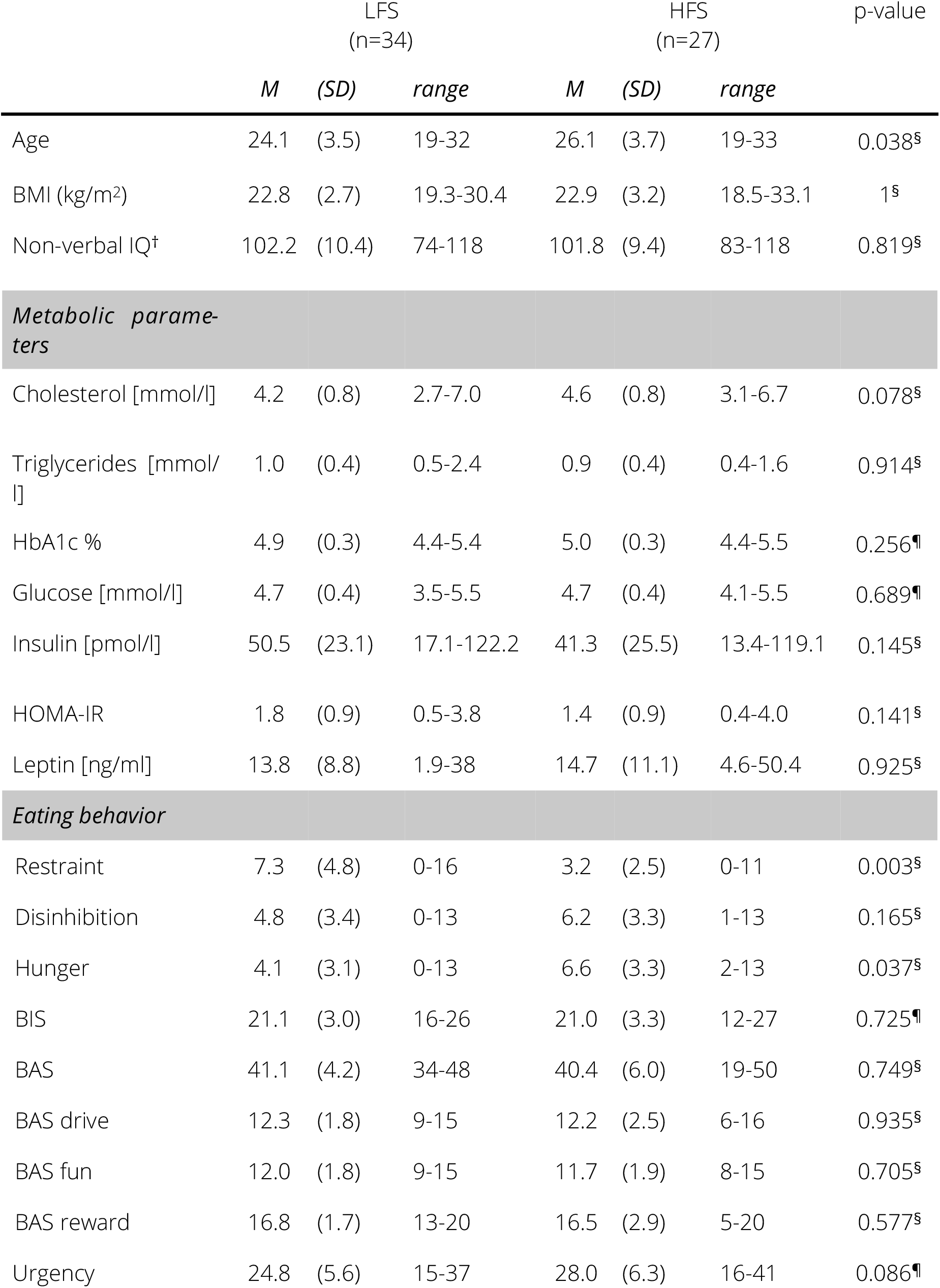
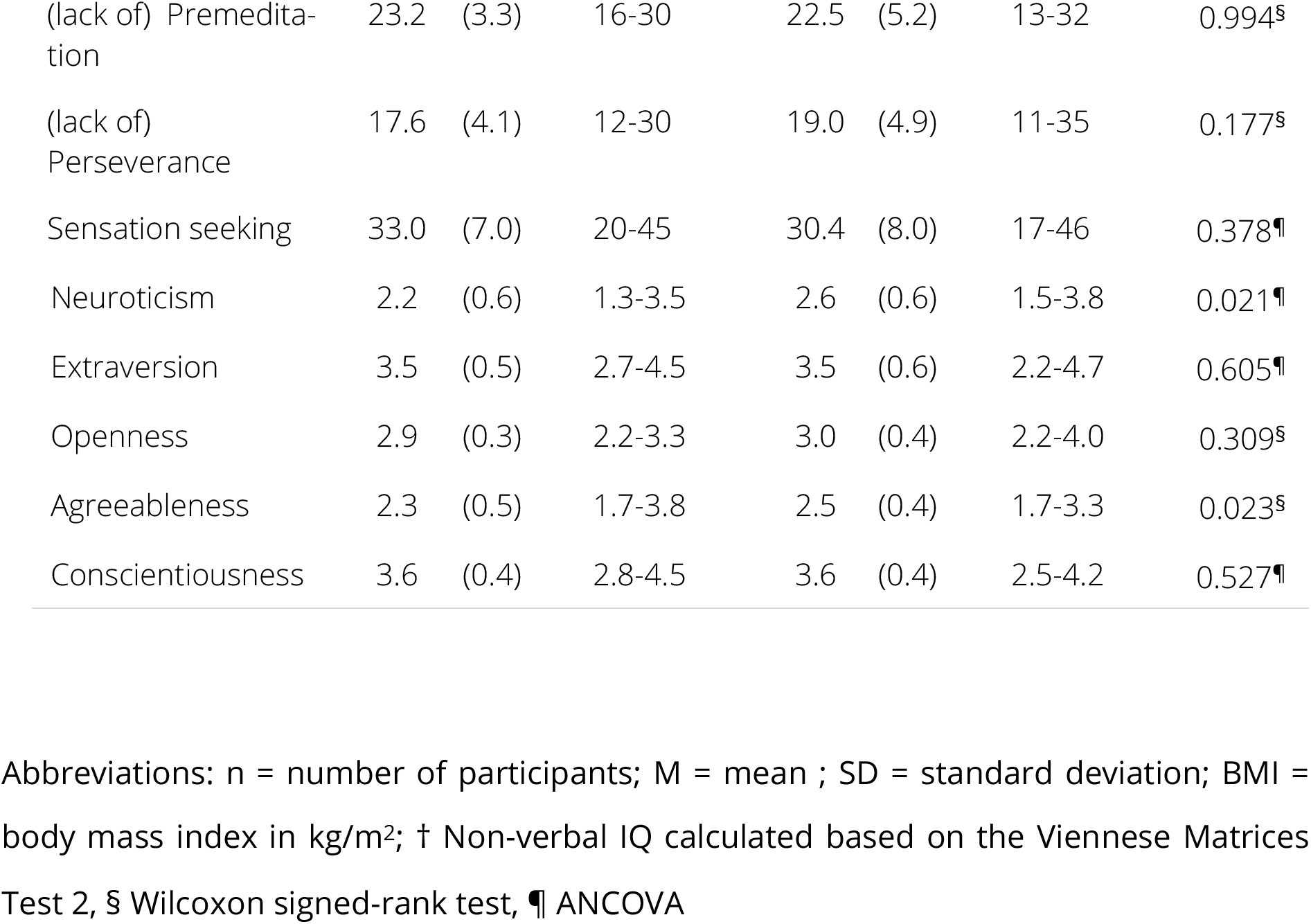
– Group demographics, metabolic parameters, eating behavior and personality traits of the extended screening sample

